# Representations of the amacrine cell population underlying retinal motion anticipation

**DOI:** 10.1101/830976

**Authors:** Michael D. Menz, Dongsoo Lee, Stephen A. Baccus

**Affiliations:** Department of Neurobiology, Stanford University School of Medicine, Stanford University

**Author notes:** Corresponding author: Stephen Baccus, 299 Campus Drive, Stanford, CA.

## Abstract

Retinal amacrine cells are a diverse population of inhibitory interneurons, posing a challenge to understand the specific roles of those interneurons in computations of the similarly diverse ganglion cell population. Here we study the predictive computation of motion anticipation, which is thought to compensate for processing delays when encoding moving objects. We recorded the membrane potential of the salamander amacrine cell population optically while recording electrically from ganglion cells with a multielectrode array. We find unexpectedly that ganglion cells with the greatest anticipation for moving stimuli exhibit a new type of predictive motion anticipation that is inconsistent with prior models of delayed inhibition. Based on the spatiotemporal correlations between thousands of amacrine and ganglion cell pairs, we modeled the contribution of the traveling wave of activity for different amacrine cell populations to the encoding of a moving bar. These models indicate that the population responses of slow biphasic amacrine cells create the greatest contribution to both types of ganglion cell motion anticipation, supporting a role for this specific amacrine cell class in the predictive encoding of moving stimuli.

**Significance Statement:** The prediction of moving stimuli is a widespread function occurring in both visual and auditory systems. The diversity of interneuron populations makes it a challenge to understand the mechanisms of these computations. To analyze how the retina anticipates motion, we optically measured inhibitory amacrine cell population activity simultaneously with electrical recording from ganglion cell populations. We then used computational modelling to assess which amacrine cell types have the right spatiotemporal responses to generate motion anticipation. In contrast to previous suggestions of a general role for inhibition, we find that slow biphasic amacrine cells specifically have the greatest contribution to motion anticipation, thus highlighting the need to directly measure and model the effects of interneuron populations in complex sensory computations.

## Introduction

Neural circuits use interneurons to transform an input neuron representation into a neural output. Understanding this process is complicated by the fact that all three neural populations carry distinct signals. Even with the advent of new recording technologies, accessing all three populations together is beyond reach in most cases. In the retina, the diversity of neural subtypes poses a significant problem towards understanding circuit function. In particular, inhibitory interneurons are extremely diverse, both in the retina (1) and higher brain (2), but most their specific functions are largely unknown. There are approximately one dozen types of bipolar cells, more than 30 amacrine cells, and more than one dozen ganglion cells. A critical barrier is the lack of the ability to observe simultaneously at multiple levels in the circuit. Furthermore, the diversity of both interneurons and retinal ganglion cells requires the observation of neural populations.

The retina produces many distinct computations involving adaptation and the detection of specific visual features (3) (4). Retinal responses to moving objects have multiple nonlinear properties, including the anticipation of moving objects (5, 6), object motion sensitivity (7) and responses to motion reversal (8). Some of these computations map specifically onto single ganglion cell types, others are spread across multiple types. Assigning those computations quantitatively to the contributions of interneurons is challenging because of the need to consider the combined effects of diverse populations of cells. Although perturbing different neural classes can potentially reveal neurons that might influence a computation, methods that can measure from diverse populations at multiple levels to evaluate which cell types are likely to generate the computation are essential to guide such perturbation experiments toward an interpretable outcome.

This study focuses on the first step in this process, the evaluation of a diverse population of cells under a specific neural computation using a computational model to assess which interneuron types are likely to generate the neural computation. We study the computation of motion anticipation, in which a moving object shifts the activated population of ganglion cells in the direction of motion, compared to the response that would be expected from a linear model of the ganglion cell response (5). Such a shift is expected to compensate for visual processing delays in the representation of a moving object. It has been proposed based on pharmacological evidence that amacrine cell inhibition generally contributes to motion anticipation (6), but the specific responses of different amacrine cells has not been evaluated as to their potential role, and there exists no computational model of the effects of the amacrine population. Here we address the question, which populations of amacrine cells have appropriate spatiotemporal responses to generate motion anticipation?

We simultaneously presented a visual stimulus to the intact retina, while recording optically from a population of interneurons using second harmonic imaging (SHG) of membrane potential (9). Simultaneously, we recorded from a population of retinal ganglion cells with a multielectrode array. This approach allows optical measurements of the membrane potential of dozens of interneurons and electrical recording from dozens of ganglion cells. From this data, we modeled the motion response in ganglion cells and in different classes of amacrine cells. Based on previous experiments injecting current into amacrine cells, we compute the expected spatiotemporal representation of the transmission from different populations of amacrine cells to individual ganglion cells.

We find that ganglion cells have two types of motion anticipation, one consistent with feed forward gain control as has previously been proposed, and a second type that anticipates motion to a greater extent that is inconsistent with previous models. By examining the responses of different amacrine cell classes, we make predictions as to the contributions of each population to the different types of motion anticipation. Based on our analysis, we predict that the largest contribution to motion anticipation of both types comes from a class of slow On-Off amacrine cell. The resulting computational model identifies a specific amacrine cell class for future, more directed single cell measurements, as well as perturbation experiments to manipulate this cell class.

## Results

### Two classes of motion anticipation in retinal ganglion cells

We performed simultaneous SHG imaging and multielectrode recording using a custom two-photon microscope in the inverted configuration (Fig. 1a, see Methods). To measure responses of amacrine and ganglion cells, we presented a smoothly moving black bar (132 µm wide, 1.26 mm/s) (5) against a light background, as the majority of ganglion cells in the salamander retina have larger Off than On responses (10). The trajectory of the bar repeated every one second.

**Figure 1.**
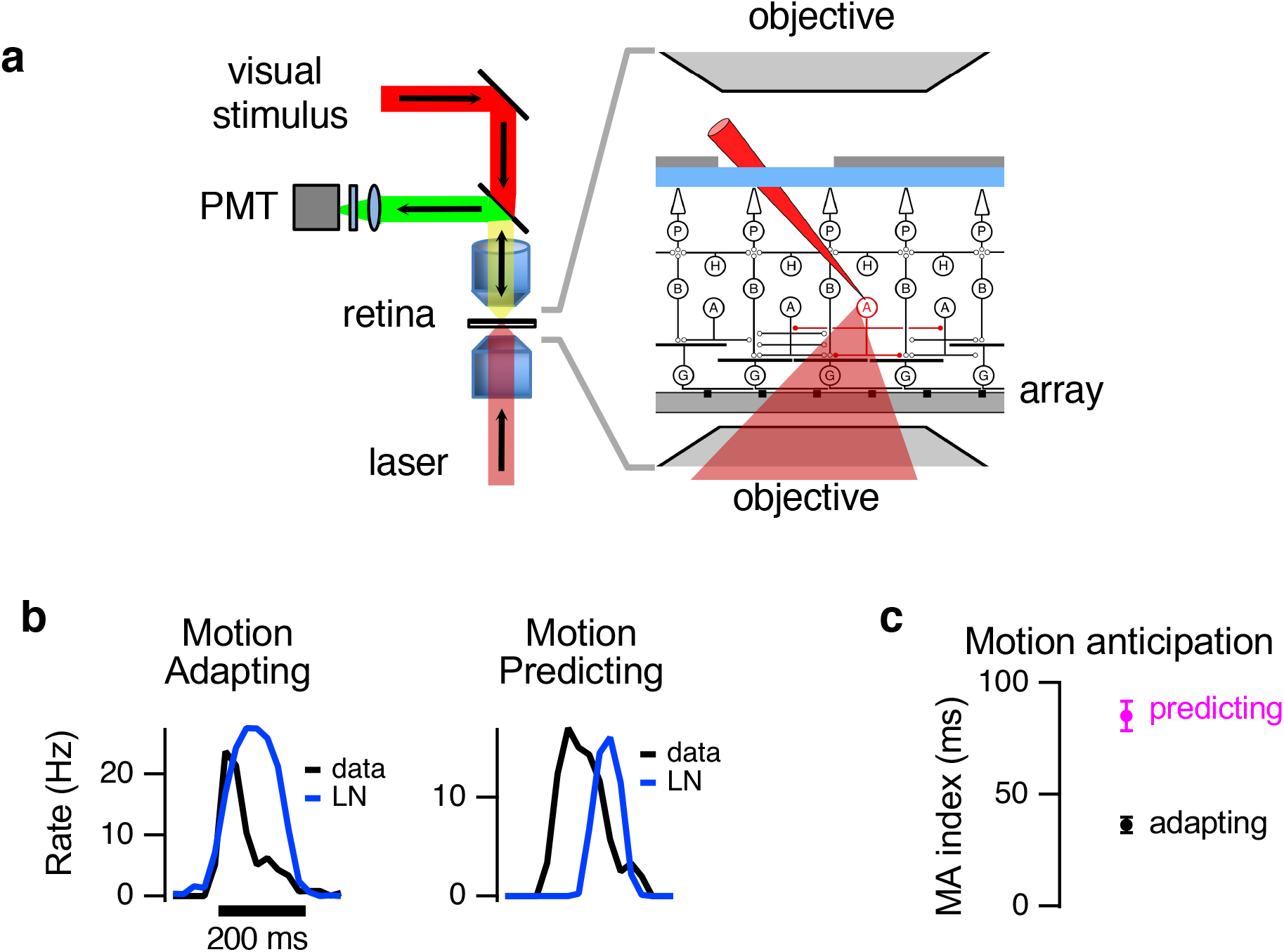
Two types of motion anticipation in the retina. (a) Setup for simultaneous two-photon imaging and multielectrode recording of visual responses. Left. Schematic diagram showing light paths for SHG illumination, collection and visual stimulation. The laser is focused through an objective below the retina. The SHG signal is collected on the opposite side of the retina by an objective and reflected to a photomultiplier tube (PMT). The upper objective also focuses the visual stimulus onto the retina. Right. Expanded view of microscope objectives and the retina, which is placed ganglion side down on a multielectrode array. Drawing is not to scale. (b) Left. An example Motion Adapting ganglion cell that shows the standard type of motion anticipation, whereby the cell’s onset of firing is at the time predicted by an LN model, but the response is truncated compared to the LN prediction. Right. An example ganglion cell in which the onset of activity substantially precedes the linear prediction, termed here a Motion Predicting cell. (c). Average motion anticipation index for Motion Predicting (mean = 85 ms) and Motion Adapting cells (mean = 36 ms) (p < 3.6e-10, Wilcoxon-Mann-Whitney two-sample rank test), defined as the temporal center of mass of firing for the data subtracted from that of the LN model.

For an object moving over a range of speeds, ganglion cell responses compensate for neural processing delays by anticipating the object’s motion, creating a traveling wave of activity that matches the leading edge of the object (5, 6). We characterized the type and extent of motion anticipation by comparing ganglion cell responses to those predicted by a simple linear-nonlinear (LN) model consisting of a spatiotemporal linear filter followed by a static (time independent) nonlinear function. We fit LN models to a white noise checkerboard stimulus, and then used the model to predict the response to a moving bar. Many cells showed motion anticipation in that the response was earlier in time on average than the LN model prediction (Fig. 1b). For these cells, in most cases the firing rate response and LN prediction began at the same time, and the cell’s response ceased earlier than the LN model’s prediction, conforming to previous models of motion anticipation that caused a truncation of the response by a reduction in gain (5, 6). We termed this type of response ‘Motion Adaptation’, referring to this decrease in gain relative to an LN model that represents a baseline non-adapting response. However, in 17 % of cells, the cell’s response began earlier than the LN model’s prediction, a type of motion anticipation that we termed ‘Motion Prediction’ (Fig. 1b). These cells exhibited greater motion anticipation than Motion Adapting cells, and Motion Predicting cells responded earlier to the moving bar than Motion Adapting cells (Fig. 1c). Motion Adapting and Motion Predicting cells did not map specifically onto ganglion cell types, as for either of the two motion types the percentage fast-Off, slow-Off and medium-Off ganglion cell classes were not substantially different (Chi squared test p > 0.3, Fig. S5D). Previous models of motion anticipation consisted of a truncation of the linear response through feedback or feedforward inhibition (5, 6). Motion Predicting cells appeared to be inconsistent with this model, and thus represented a second type of motion anticipation. We test this idea further below.

### Visual responses to a moving bar in amacrine and ganglion cell populations

To identify amacrine populations that could potentially deliver the appropriate spatiotemporal input to account for motion anticipation, we imaged in the inner nuclear layer immediately proximal (within ∼5 microns) of the boundary of the inner plexiform layer (IPL) (Fig. 2a,b, Fig S1-3). Although the inner nuclear layer overall also contains bipolar and horizontal cells, this section immediately proximal to the IPL is exclusively amacrine cells as determined by previous immunostaining and is termed the amacrine cell layer (6-8).

**Figure 2.**
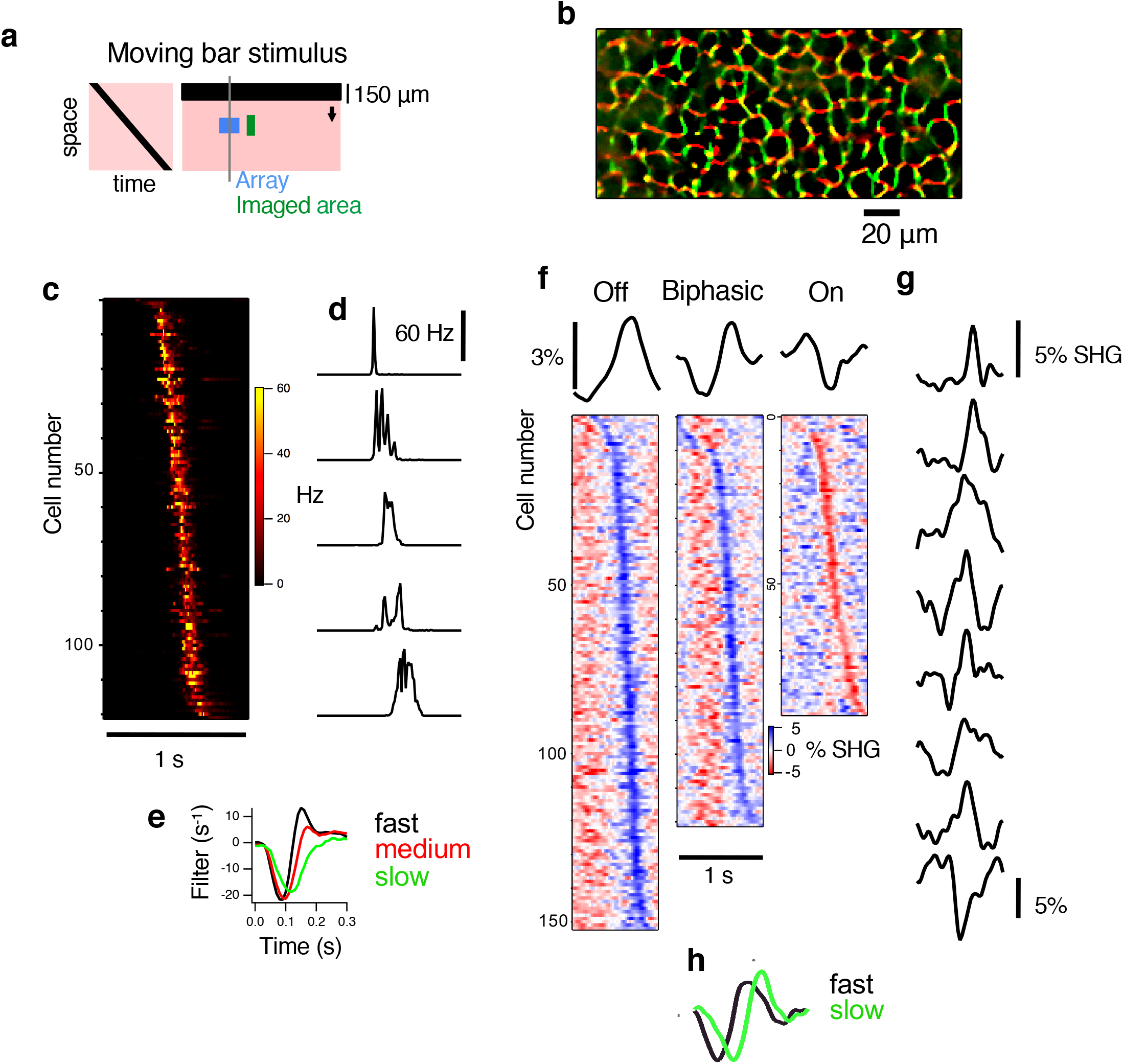
Simultaneous recording of amacrine and ganglion cell populations to moving stimuli. (a) Left. Space-time plot of the trajectory of the bar shown at a spatial cross section indicated by the vertical grey line at right. Right. Spatial layout of visual stimulus, electrode arrays and imaged areas as projected onto the retina. A 132 μm black bar moved at a speed of 1.26 mm/s against a bright background (pink area), which was colored red to avoid crosstalk with the SHG emission signal. The electrode array had two groups of 30 electrodes each covering an area of 150 μm x 120 μm and spaced 620 μm apart. Four locations were imaged sequentially in a single retina. Each location was laterally offset (88 μm from the border of the array to the border of the imaged area) to an electrode group. The electrode array is indicated by the blue rectangle. The green rectangle indicates one area imaged using the two-photon microscope. (b) An SHG image in the inner nuclear layer just above the inner plexiform layer, 55 *µ*m above the array. Two recordings were made 90° apart in polarization angle and combined in two different color channels of the image, with 0° in green and 90° in red, indicating strong polarization dependence as expected from the SHG signal. Yellow pixels respond equally to both polarization angles. (c) Left. Peristimulus time histograms (PSTHs) of 122 ganglion cell PSTHs ordered vertically by latency, pooled over four retinas. Each row is a different cell, and color indicates firing rate. (d) Example PSTHs from five ganglion cells. (e) Ganglion cells were clustered by k-means into fast, medium and slow types based on temporal filters measured using a white noise checkerboard stimulus. (f) SHG motion responses from 365 amacrine cells pooled over four retinas categorized into three classes, Off, Biphasic and On. Responses are ordered by latency. (g) Example SHG responses are shown for eight cells. (h) Biphasic amacrine cells are further subdivided into fast and slow by k-means clustering to be used in the model of Fig. 4.

In response to a uniform field 1 Hz flash or a moving bar presented at 1 Hz, amacrine cells showed either Transient Off, Transient On, On-Off, Sustained Off, and Sustained On responses (Fig. S4). These responses appeared qualitatively similar to intracellular electrical recordings performed in separate retinas.

In four retinas, we recorded 122 ganglion cells, 365 amacrine cells and 4812 amacrine-ganglion simultaneously recorded cell pairs. Amacrine and ganglion cells responded at different times depending on their spatial position (Fig. 2 c-f), with Off-type ganglion cells responding to the moving bar with short bursts of spikes occurring at different times and durations. Off-type ganglion cells were classified into “slow”, “medium”, and “fast” based on their temporal filters measured from a checkerboard white noise stimulus (Fig 2d).

Amacrine cells were classified at first into three types depending on their response to the dark moving bar: depolarizing (Off type, 153 cells), hyperpolarizing followed by depolarizing (biphasic type, 122 cells) or hyperpolarizing (On type, 90 cells), with latencies and durations that differed between cells (Fig. 2, e-g). Intracellular responses to the moving bar did not necessarily correspond in a one to one manner to the flash response, however cells having similar responses to the optically recorded biphasic type were On-Off cells (Fig S4b).

### The neural image of the amacrine population at the time of a ganglion cell spike

For a moving bar stimulus, we computed the average amacrine cell response triggered on the spike of the ganglion cell as a function of spatial distance from the ganglion cell and temporal delay. Figure 3 shows examples of amacrine cell responses relative to ganglion cell spiking for On, Biphasic and Off amacrine cells. By combining many cell pairs at different locations, we constructed a spatiotemporal representation of the average Off amacrine cell response at different distances from a Fast-Off-type ganglion cell and at different times relative to the that ganglion cell’s spike (Fig. 3b). This map showed a wave of amacrine cell activity sweeping across the retina. The velocity of the peak of the wave was 1.27 mm/s, matching closely the actual bar velocity of 1.26 mm/s.

**Figure 3.**
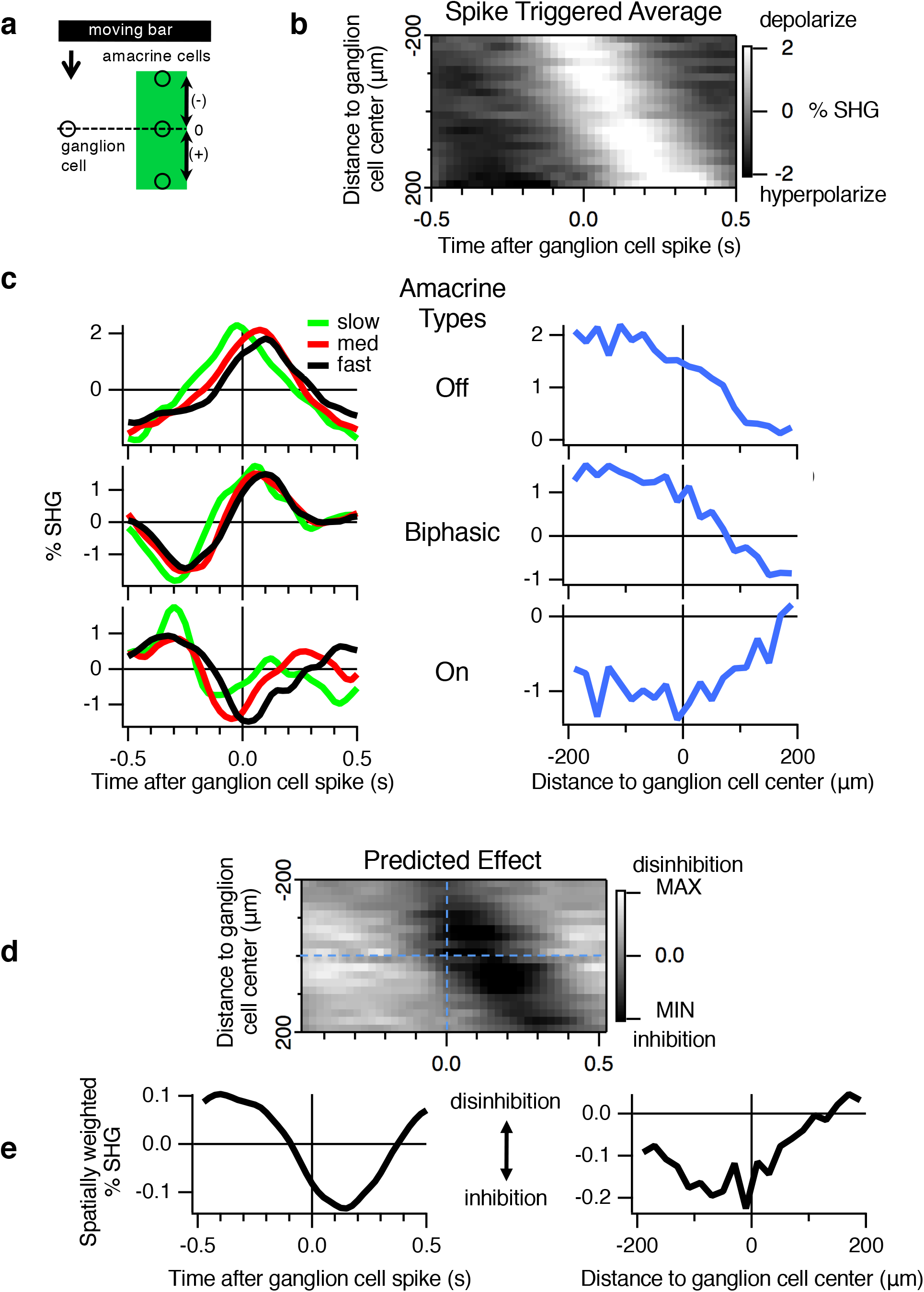
Prediction of effects of the amacrine cell population on ganglion cell responses. (a) Diagram depicting the distance between amacrine and ganglion cells measured along the direction of a moving bar. (b) Space-time plot of the average population Off amacrine activity relative to the time of a spike of Fast Off ganglion cells (n=835 pairs), computed as the spike-triggered average of amacrine cell responses binned by amacrine-ganglion cell distance in the direction of the moving bar. Cellular location was taken as the amacrine soma location and the ganglion cell receptive field center. Negative values of stimulus distance indicate the bar encountered the amacrine cell before the ganglion. (c) Left. Temporal slices of amacrine-ganglion cell correlations for different amacrine and ganglion cell classes for nearby cell pairs (< 20 µm in the direction of motion). Top Left. Temporal slices from space-time plots taken with Off amacrine cells and Off ganglion cells (fast, 835 pairs, medium, 965 pairs, slow, 272 pairs). Middle Left. Temporal slices from space-time plots taken with biphasic amacrine and Off ganglion cells (fast, 692 pairs, medium, 715 pairs, slow, 189 pairs). Bottom Left. Temporal slices from space-time plots taken with On amacrines and Off ganglion cells (fast, 472 pairs, medium, 484 pairs, slow, 169 pairs). Right. Spatial slices from space-time plots of different amacrine cell types averaged across all ganglion cells presented when the ganglion cell spikes. Top Right. Spatial slice of space-time plot for Off amacrines averaged across all ganglion cells (n=2090 total pairs). Middle Right. Spatial slice of space-time plot for Biphasic amacrines averaged across all ganglion cells (1596 total pairs). Bottom Right. Spatial slice of space-time plot for On amacrines averaged across all ganglion cells (1125 total pairs). (d) The spike-triggered average responses from panel (b) were filtered by an average amacrine cell transmission filter (25 ms delay) computed from previous intracellular studies (13) and spatially weighted with a space constant of 83 microns (13), then inverted in sign to predict the average effect the Off amacrine cell population on the average Fast Off ganglion cell, relative disinhibition in white, inhibition in black. (e) Left. Integration along the spatial axis from of the prediction from panel (d) yields an estimate of the relative disinhibition and inhibition an average ‘fast’ off ganglion cell would receive from a spatially distributed population of Off amacrine cells. Right. Spatial distribution of inhibition from panel (d) taken within 25 ms of a ganglion cell spike.

To examine cell type differences in the population response, we constructed such a space-time map for each combination of ganglion cell type (fast, medium, slow) and each amacrine cell type (Off, Biphasic and On). Fig. 3c shows the temporal distribution of amacrine responses for nearby (within 20 µm) amacrine-ganglion cell pairs for all nine combinations of cell pair types. In general, for amacrine cells in the same spatial location, the peak amacrine depolarization occurred soon after the ganglion cell spiked for Off and Biphasic type amacrine cells (Fig. 3c), with variations in timing that are dependent on ganglion cell type. For On amacrine cells, which are expected to provide net excitation to Off-type ganglion cells through disinhibition (11, 12), peak hyperpolarization occurs near the time of the ganglion cell spike, with membrane potential reaching a minimum either just before or after the ganglion cell spike depending on the ganglion cell type. Examining the spatial distribution of the amacrine cell response at the time ganglion cell’s spike, we found that the most depolarized amacrine cells trail the moving bar.

We then used the spatiotemporal map of amacrine activity to predict the effect that each Off amacrine cells would have on Fast-Off ganglion cells (Fig 3d). To do so, we needed to estimate the timing and spatial extent and temporal delay of their transmission. For sustained Off-type amacrine cells, we used previous measurements of their temporal transmission filter computed from intracellular studies, which we modeled as an exponential spatial decay of 83 µm, and a monophasic negative temporal filter with a time to peak of 25 ms (13). After transforming the amacrine population response with this transmission filter, the result is a spatiotemporal map of the estimated effect from amacrine cells at each spatial position and time. Integrating across space yielded an estimate of the total effect of the amacrine population as a function of time relative to a ganglion cell’s spike (Fig 3e Left). From this estimated population effect, maximum disinhibition occurs several hundred millisecond before spiking, inhibition during spiking, and maximum inhibition at about 150 ms after spiking. Note that in this case, disinhibition refers to a decrease in inhibition below a relative baseline. Then at the time of a ganglion cell spike, we examined the spatial distribution of amacrine effects (Fig 3e, Right). The largest inhibition is coming from nearby cells in the direction the bar has just passed (left of zero), which is a consequence of the spatial weighting. The distribution around zero was not symmetrical, with more inhibition coming from the amacrine cells that preceded the ganglion cells.

Comparing Biphasic and Off-type ganglion cells, although there are differences in the timing of individual amacrine cells, if one averages across all individual amacrines of one type (Fig. 3c), both cell types depolarize just after the ganglion cell spike, which would be expected to truncate ganglion cell activity consistent with motion anticipation. However, Biphasic and Off-type ganglion cells had a different timing of hyperpolarization before the ganglion cell spike, with Biphasic cells hyperpolarizing just before the ganglion cell spike.

### Specific amacrine types create different inputs during motion anticipation

To test the potential role of different amacrine cell populations in motion anticipation for these two classes of ganglion cells, we used the measured amacrine cell population responses to account for the difference in the ganglion cell response from an LN model. We clustered the amacrine responses into four types, based on their responses to moving bars, as shown in Fig 2e (Off, Fast Biphasic, Slow Biphasic and On). These responses were then delayed in time by 25 ms and filtered spatially as in Fig. 4 to represent the effect of each amacrine cell on each ganglion cell.

**Figure 4.**
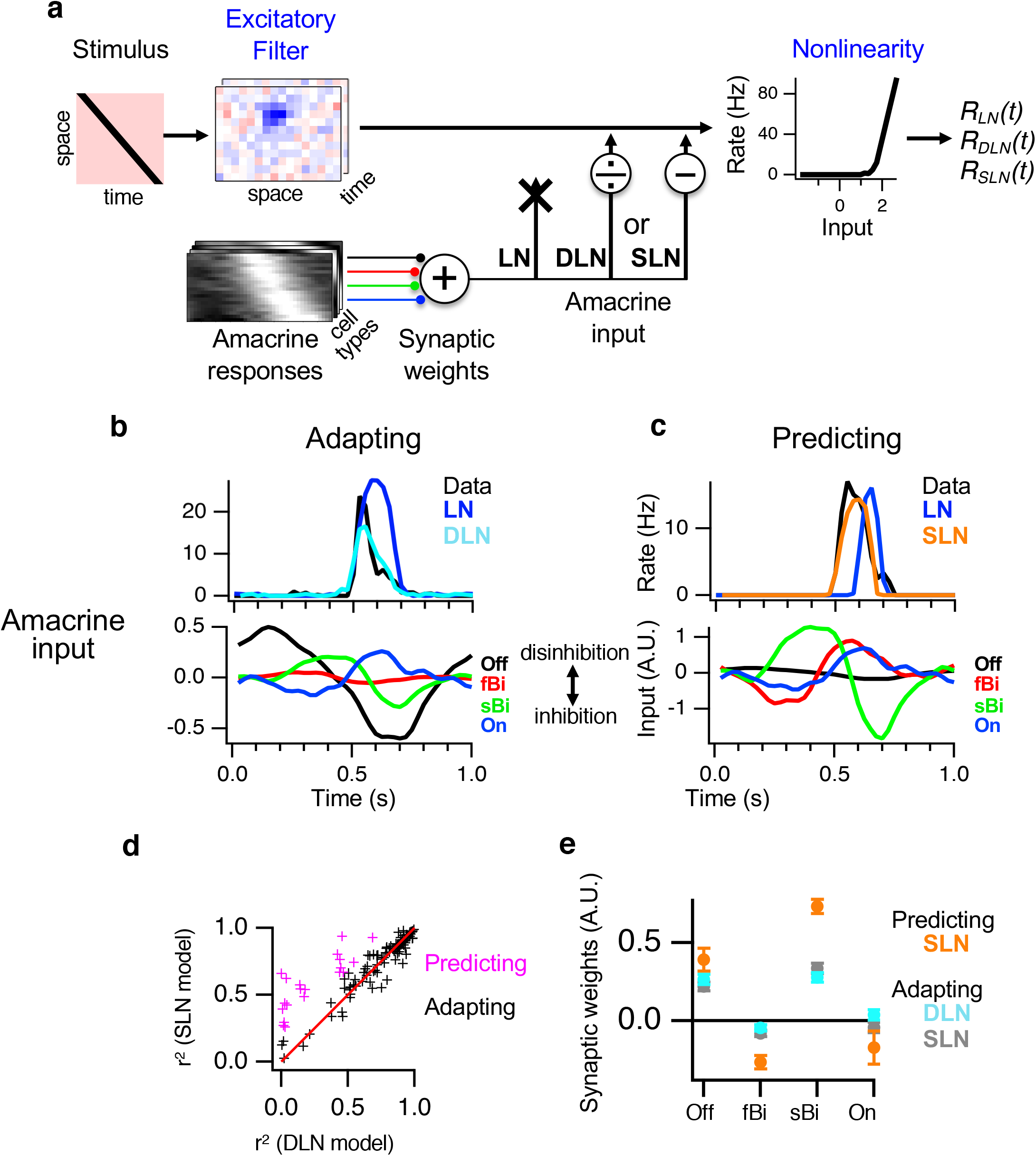
Specific amacrine responses underlie motion anticipation. (a) Models of motion anticipation combining the linear receptive field with measured amacrine cell responses. The moving bar stimulus S(x,t) is convolved with the space-time receptive field F(x,y,t) of the ganglion cell as measured with white noise. This linear prediction serves as the first stage of one of three models 1) A Linear-Nonlinear (LN) model that simply passes the linear prediction through a threshold. 2) The gain of the linear prediction is divisively controlled by a weighted linear combination of amacrine cell responses prior to the nonlinearity (Divisive LN or DLN model) 3). The linear prediction is summed with a weighted linear combination of amacrine cell responses prior to the nonlinearity (Subtractive LN or SLN model). Four scalar weights on different amacrine cell classes are optimized to fit the model’s output to the data (one for each of Off, slow Biphasic, fast Biphasic and On amacrine cell populations). (b) Top. An example of a Motion Adapting cell ganglion cell with a motion anticipation index of 26 ms (difference in the temporal center of mass of firing rate, LN vs. Data), along with the prediction of the DLN model. Bottom. For the same cell, the weighted amacrine cell input averaged over the total amacrine population for four different amacrine cell classes, using the optimal weighting for the DLN model (one scalar weight for each of four cell classes). Traces are shown inverted so that inhibition is down, disinhibition is up. (c) Top. An example Motion Predicting ganglion cell with a motion anticipation index of 60 ms, along with the prediction of the SLN model. Bottom. Weighted amacrine input, again shown with inhibition in the downward direction. (d) Scatterplot of the squared correlation coefficient (r^2^) for the SLN model (subtractive) vs. data compared to the DLN model (divisive). Red line is the identity line. Cells were classified as Motion Adapting or Motion Predicting by whether the SLN fit exceeded the DLN fit (Fig. S5b). (e) Average amacrine cell synaptic weights by model type (DLN or SLN) and ganglion cell type (Adapting or Predicting) and amacrine type (Off, fast Biphasic, slow Biphasic, and On). The DLN model did not effectively fit Motion Predicting ganglion cells.

We first considered as previously proposed whether a reduction in gain of the ganglion cell with respect to its linear input could produce motion anticipation. We tested this idea by modeling whether amacrine responses could act to control the gain of ganglion cells via a divisive mechanism. This Divisive Linear Nonlinear (DLN) was a modification of the LN model that took the measured amacrine responses and controlled the gain of the output of the linear filter via a divisive effect. This model took the same parameters as the LN model, but included four additional parameters that represented the weighting of each of the four amacrine cell classes of the optically measured responses. Because we began with an LN model fit to white noise without using the amacrine cell recordings, the model did not include any contribution of amacrine cells to this initial linear filter, although one that expects that such a contribution does occur. Thus, in the model the amacrine contribution represents strictly the nonlinear effects of these cells that occur for a moving stimulus that differ from effects during white noise. The DLN model successfully fit Motion Adapting cells by causing a decrease in gain that caused the firing rate to drop to zero earlier than predicted by the LN model. However, the DLN model could not capture the response of Motion Predicting ganglion cells. This was because compared to the prediction of the LN model, Motion Predicting ganglion cells had an earlier onset of firing. Therefore, a gain change that change the amplitude of firing could not shift the response to achieve this earlier onset. This result confirmed that Motion Prediction represented a distinct type of motion anticipation that was inconsistent with prior models, where motion simply cause a delayed gain change with respect to the linear visual input.

We then tested a Subtractive Linear Nonlinear (SLN) model that subtracted amacrine responses from the linear filter output of the LN model, again with one weight parameter for each of the four amacrine classes. In addition to capturing the response of Motion Adapting ganglion cells, the SLN model could also fit Motion Predicting ganglion cells. The difference in the fits of the SLN and DLN models were used as an additional classification as to whether a cell exhibited Motion Adaptation or Motion Prediction (Fig. 4d, S5b).

To assess whether certain amacrine cell types had the appropriate spatiotemporal responses to potentially contribute to motion anticipation, and to test the idea that nonspecific inhibition contributed motion anticipation, we examined the weights of the different amacrine cell classes. We found that the weights in the model were different for different amacrine types. For Motion Adapting cells, both DLN and SLN models had similar values, with slow biphasic and Off amacrine cells contributing the most, and On type amacrine cells contributing little (Fig. 4e). For the SLN model of Motion Predicting cells, slow and fast biphasic amacrine cells had the greatest contribution. Weights were allowed to be of either sign because of the potential of polysynaptic disinhibitory connections, and fast biphasic amacrine cells had a negative weight, which corresponded to a disinhibitory effect. In addition, across the population, the weights of slow biphasic amacrine cells were strongly correlated with the amount of motion anticipation across cells, with cells that showed greater motion anticipation having a greater weight in the model from slow biphasic cells (r = 0.7, Fig. S5c).

By examining the relative timing of amacrine cells, ganglion cell linear predictions and the ganglion cell response, we can infer how the different types of motion anticipation arise. In response to a moving dark bar, biphasic amacrine cells hyperpolarize then depolarize, creating disinhibition followed by inhibition. If the time interval of disinhibition overlaps the leading edge of excitation, then disinhibition will shift the onset of ganglion cell firing earlier, leading to the Motion Prediction response (Fig. 5). However, if the time interval of disinhibition is earlier compared to excitation, then disinhibition does not act at the correct time to combine with subthreshold excitation. Instead, the only effect is the truncation of activity by inhibition, leading to Motion Adaptation. Although Motion Predicting and Motion Adapting cells both received input from Off type and slow biphasic amacrine cells, Motion Predicting cells had a greater contribution of slow biphasic amacrine cells, which created larger disinhibition near the onset of ganglion cell firing. These results provide a testable quantitative model whereby slow On-Off amacrine cells generate Motion Prediction and contribute to Motion Adaptation, and both Off and slow On-Off amacrine cells contribute to Motion Adaptation.

**Figure 5.**
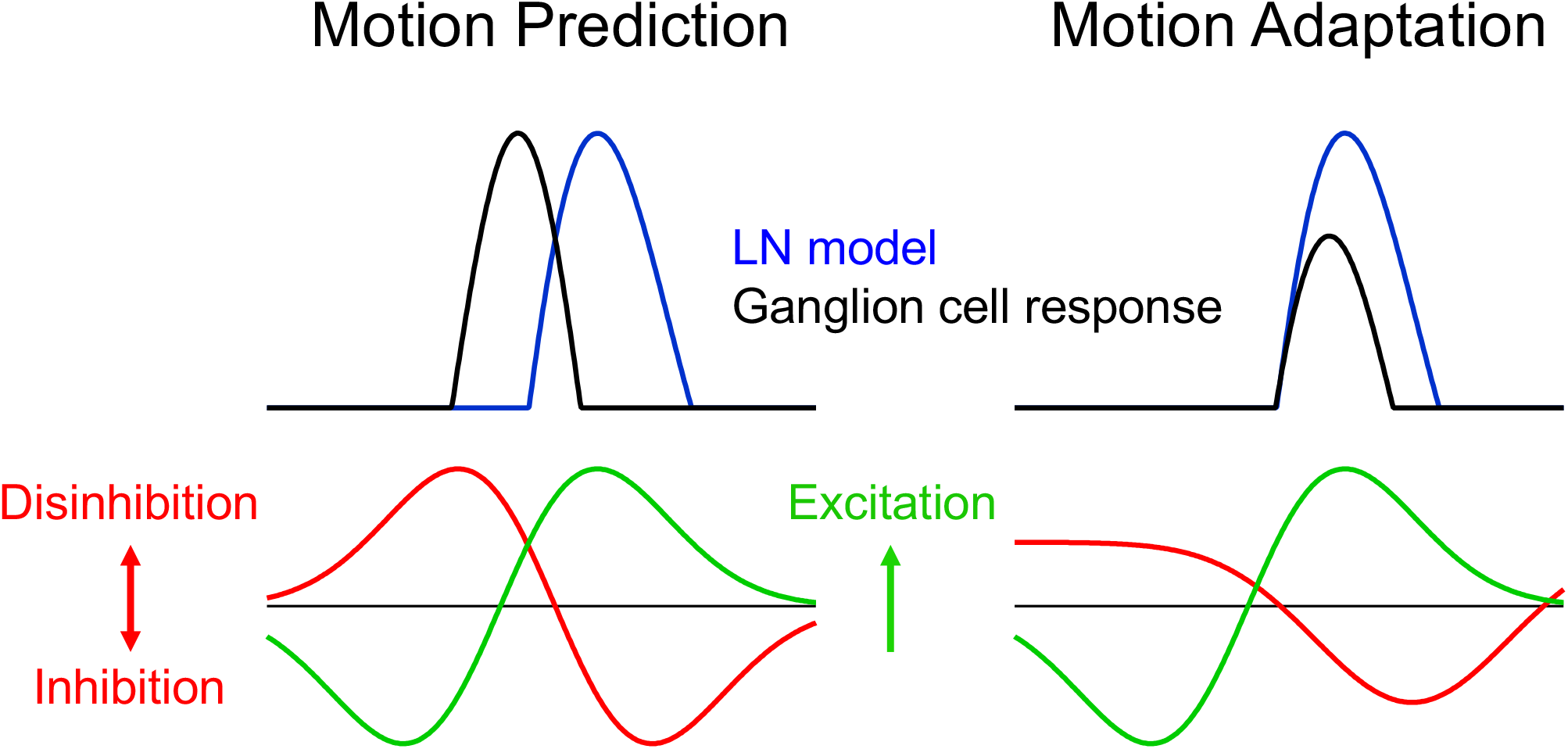
Model of amacrine contribution to motion anticipation. Left. Schematic summary of model depicting Motion Prediction. Disinhibition from slow biphasic amacrine cell hyperpolarization arrives during the leading edge of excitation from bipolar cells, thereby advancing the onset if firing. In addition, delayed inhibition truncates activity. Right. For Motion Adaptation, disinhibition from sustained Off-type amacrine cells does not arrive near the leading edge of excitation, and only delayed inhibition has an effect.

## Discussion

In this study we have controlled visual input to a population of photoreceptors, while simultaneously measuring optically from a population of interneurons and recording electrically from a population of ganglion cells. In doing so, we have examined populations of amacrine cells to assess which amacrine types have spatiotemporal responses that make them likely to contribute to motion anticipation, and to determine how the computation is synthesized. Previously it was proposed that motion anticipation results from general inhibitory properties of the retina (Johnston and Lagnado, 2015). In contrast, our results point to distinct, specific populations of amacrine cells as contributing to motion anticipation. Specifically, On-Off amacrine cells with slow biphasic responses to a moving bar are the most likely cell classes to contribute to motion anticipation, in particular for Motion Predicting ganglion cells. The relative timing of amacrine cell disinhibition and inhibition compared to ganglion cell excitation appears to play a critical role in determining the type and magnitude of motion anticipation.

It has been previously suggested that a gain control mechanism could generate motion anticipation (5). We confirm that such a mechanism (our divisive model) can account for the motion anticipation for most ganglion cells. However, it is clear that Motion Predicting cells have a response that could not be generated simply by a divisive reduction in gain applied to the linear response (Fig 5). A subtractive effect of amacrine inhibition is intrinsically more versatile because the ganglion cell membrane voltage can be depolarized or hyperpolarized at any time interval.

Our examination of the relative timing of amacrine and ganglion cell responses highlight the need to consider the detailed spatiotemporal responses of interneuron populations in proposing circuit explanations for neural computations. Previous studies perturbing sustained Off amacrine cells with direct current injection showed that the dominant action of these amacrine cells was through hyperpolarization so as to create disinhibition, thereby effectively causing net excitation of Off-type ganglion cells (14). These previous studies however, were for spatiotemporal white noise, and our studies here indicate that for a moving bar stimulus disinhibition from Off type amacrine cells does not arrive at the proper time to shift the onset of ganglion cell firing earlier in time in the case of Motion Predicting cells. Further studies making similar timed perturbations during moving stimuli will be needed to directly test the models proposed here.

One might look at the structure of the SLN model and note that if the amacrine response was itself linear, then the SLN model would be, in fact, an LN model that linearly sums amacrine and non-amacrine input followed by a threshold. But this would be a contradiction to the observation that an LN model fit to white noise does not fit the ganglion cell response to motion. Thus, the failure of the LN model fit to white noise, but the success of the SLN model for motion implies that either the amacrine cell response or amacrine cell transmission has a nonlinear response to motion compared to white noise.

It has also been proposed that amacrine cell inhibition could be the mechanism responsible for gain control (6) and we confirm that as well. However, this previous work suggested a very simple model that consists of an excess of inhibitory inputs over excitatory inputs that are both randomly distributed. Our study is more consistent with a circuit that has specific connections between specific cell types as is known for other motion-related phenomena such as direction selectivity and objection motion sensitivity (15) (16). Comparing Motion Predicting to Motion Adaptive cells in the subtractive model, the greatest difference between the two cell classes is the value of the weights from slow Biphasic amacrines (Fig 5e). Furthermore, slow Biphasic amacrines have the strongest correlation with motion anticipation (Fig. S3c). These results suggest that slow Biphasic amacrines play a more important role in motion anticipation compared to the other amacrine cell types, in particular as to cells that exhibit the greatest amount of motion anticipation, the Motion Predicting cells. For Motion Adapting cells, Off and slow Biphasic amacrine cells make a similar contribution.

The models that include measured amacrine cell responses account for the deviation in ganglion cell response from the expected output of the ganglion cell linear receptive field, a deviation that represents the motion anticipation phenomenon (5, 6). But it may well be that the amacrine cells also contribute to that linear receptive field, and that amacrine cell linear contribution is not explicitly shown in our models. A more mechanistic model of motion anticipation would eliminate this distinction between the amacrine cell contribution to the linear receptive field and nonlinear response, and just define one contribution of amacrine cells that accounts for both. This model would require optical recording of the amacrine population under a more rich stimulus set such as white noise to map receptive fields, which we have not done here. To complete such models, direct perturbation of individual amacrine cell types (Manu & Baccus, 2011; Kastner et al., 2019) will be needed to define their precise spatiotemporal contributions to motion anticipation. Our population measurements from interneurons and retinal outputs will direct such future studies, and point out the benefit of simultaneous recording of interneurons and circuit output to understand how diverse types of interneurons shape the diverse types of ganglion cells in the retina.

## Acknowledgments

This work was supported by grants from the Pew Charitable Trusts, McKnight Endowment Fund for Neuroscience, the Weston Havens Foundation, the Alfred P. Sloan Foundation and the E. Matilda Ziegler Foundation (S.A.B.). We thank Mihai Manu and David Kastner for intracellular recording of amacrine cell responses.

## Supplemental Figure Legends

**Figure S1.**
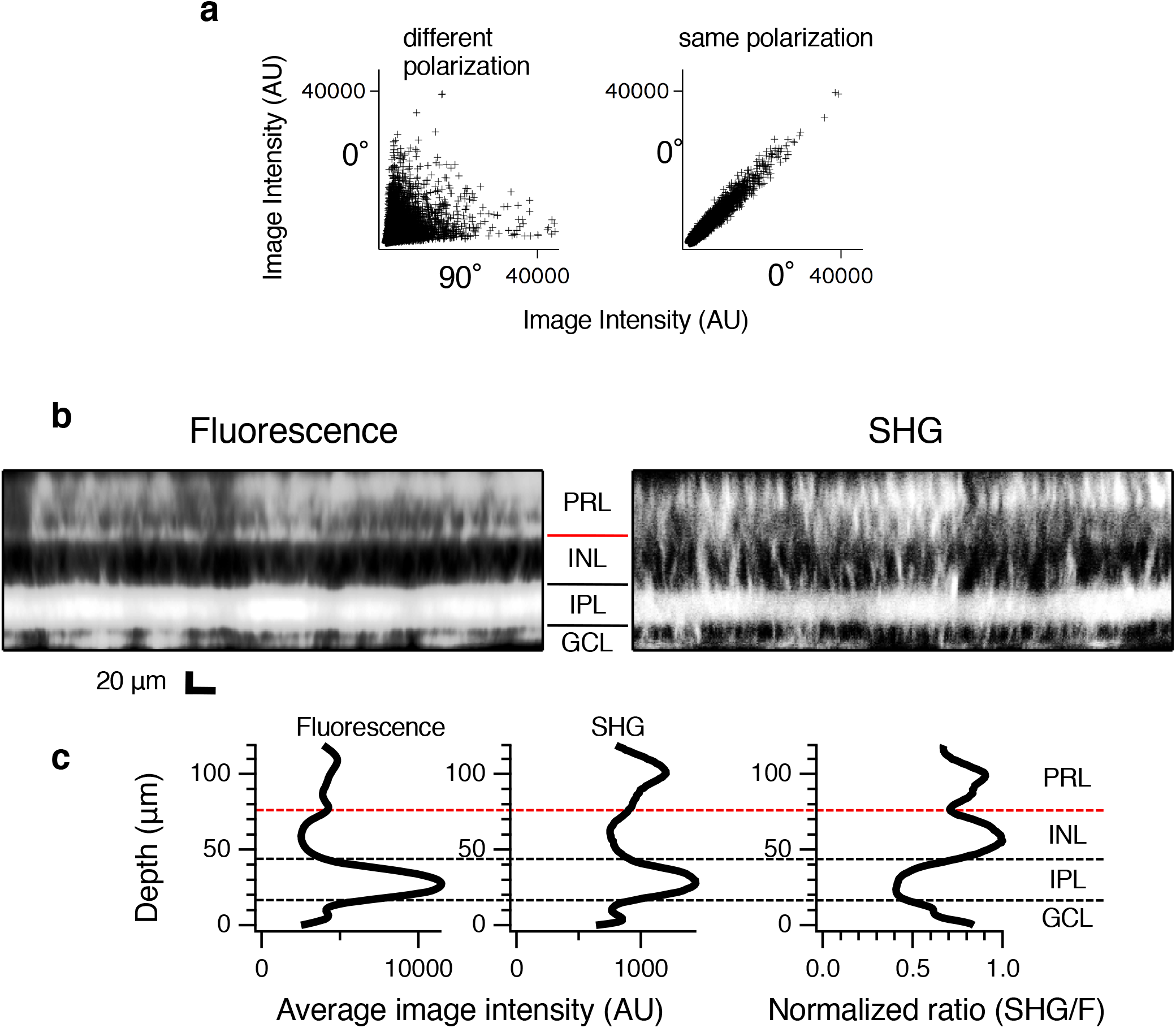
Second harmonic generation imaging of the retina. (a) Scatterplot of all pixel values from the SHG image in Fig. 2b for different polarization angles (left) or from two separate images at the same polarization angle (right). (b) Cross-section (XZ) projection of retinal stack, with the array at bottom. Left. Fluorescent image. PRL=photoreceptor layer, red line is outer plexiform layer, INL=inner nuclear layer, IPL=inner plexiform layer, GCL=ganglion cell layer. Right. Second Harmonic Signal (SHG) image (c) Left. Depth (Z) projection of fluorescent image from (b). Middle. Depth (Z) projection of SHG image from (b). Right. Normalized ratio of SHG to Fluorescence depth projections. Dotted lines indicate borders between layers, red dotted line indicates center of outer plexiform layer.

**Figure S2.**
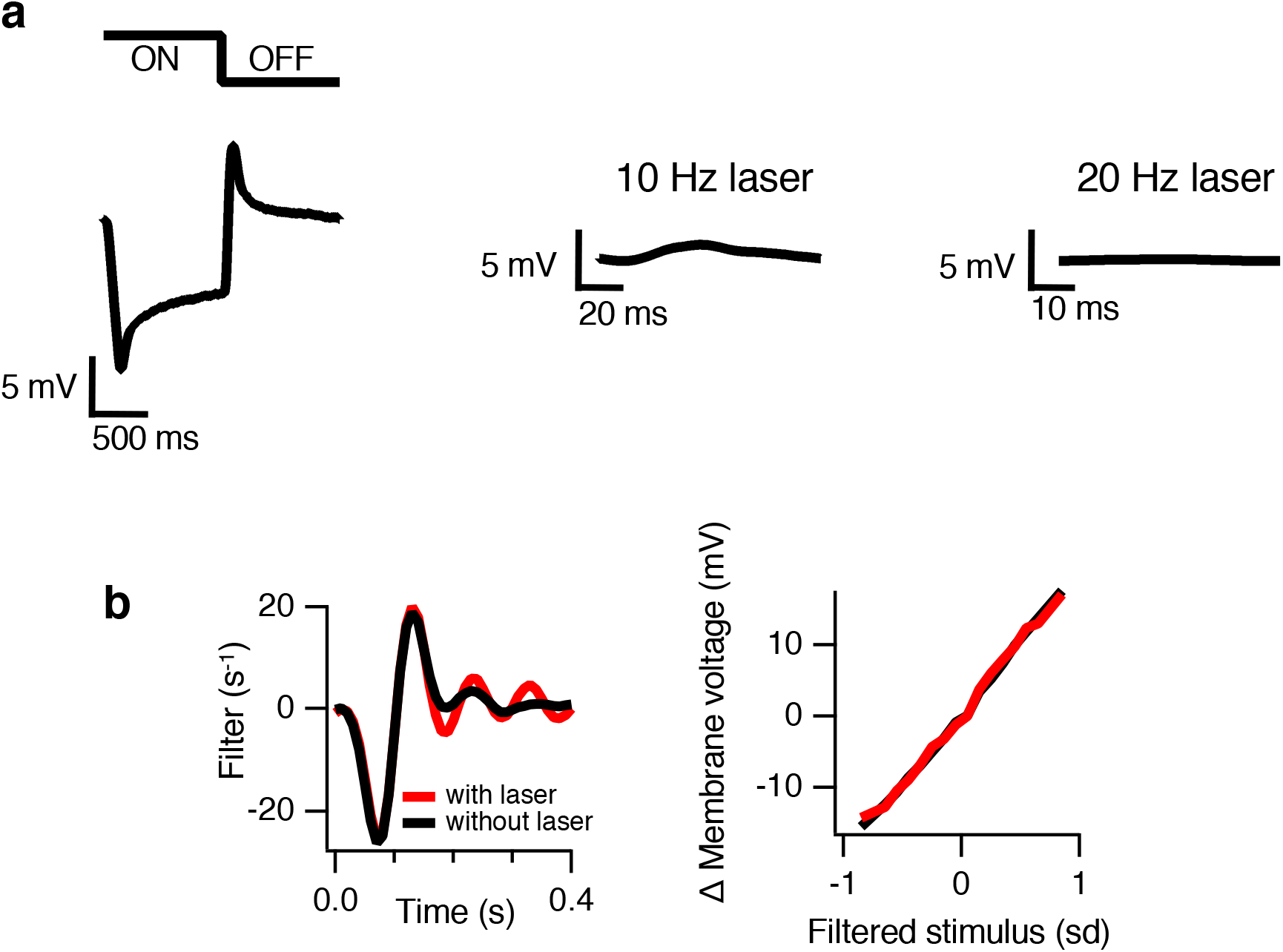
Intracellular recording of a sustained Off amacrine cell with and without laser scanning. (a) Left. Average response to 0.5 Hz uniform flash. Middle. Average response to laser scanning at 10Hz. Right. Average response to laser scanning at 20Hz. (b) LN model measured with uniform gaussian white noise stimulus with 12% contrast without laser scanning (black) and with 10 Hz laser scanning (red). Left, temporal filter. Right. Nonlinearity of LN model, which is linear in this case, showing that the visual response gain does not change when the laser is on.

**Figure S3.**
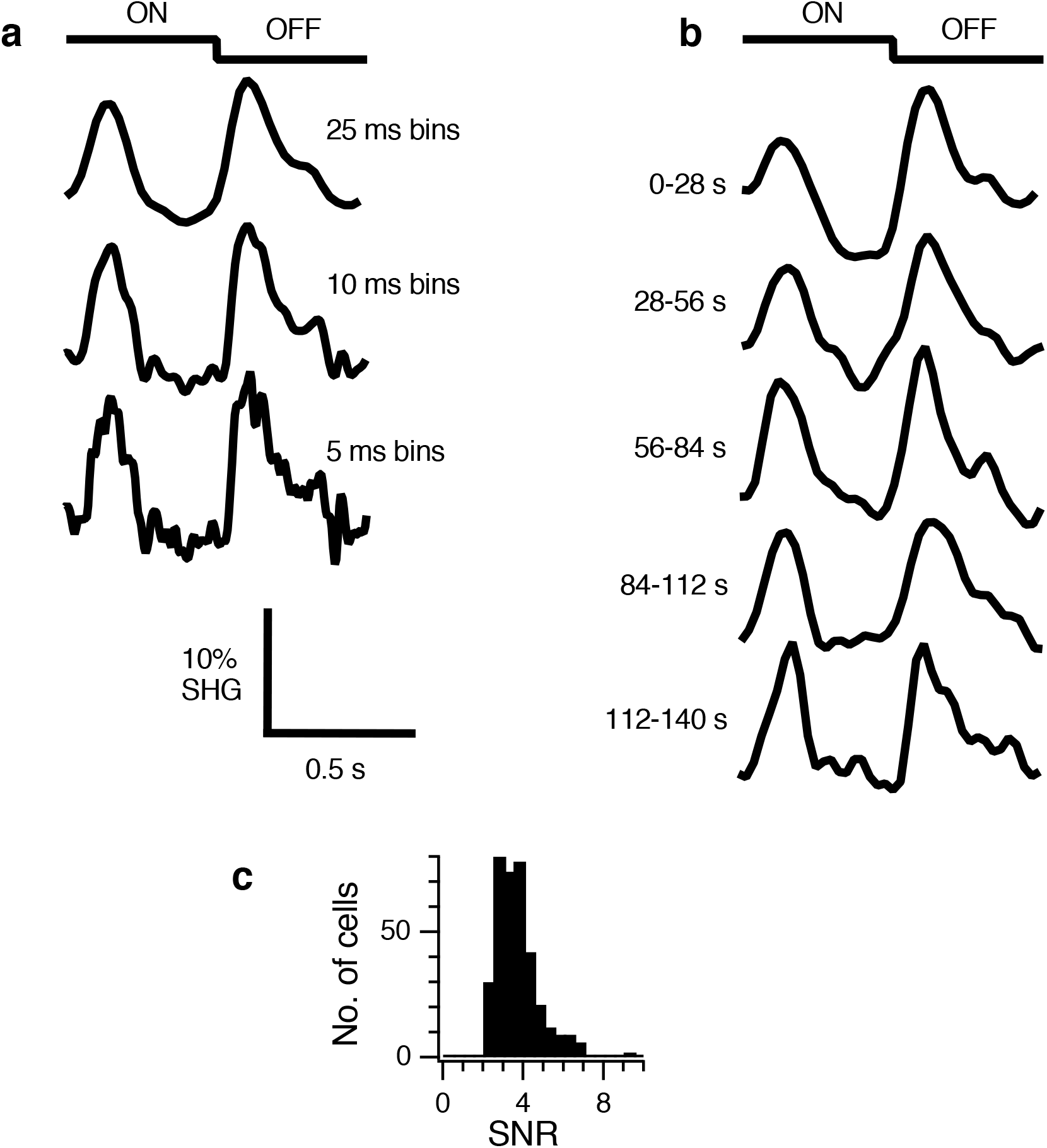
Signal-to-Noise of SHG recordings. (a) ROI intensity from one amacrine cell binned relative to stimulus time (uniform field, one second on, one second off). Top. 25 ms bins. Middle. 10 ms bins. Bottom. 5 ms bins. (b) Same cell as in (a), binned at 25 ms, with the total 140 seconds of recording time subdivided into five equal parts showing how the average response changes during the recording. (c) Histogram of signal-to-noise ratio for 365 amacrine cells (average is 3.7). A value of 2.0 was the minimum to be included in the study.

**Figure S4.**
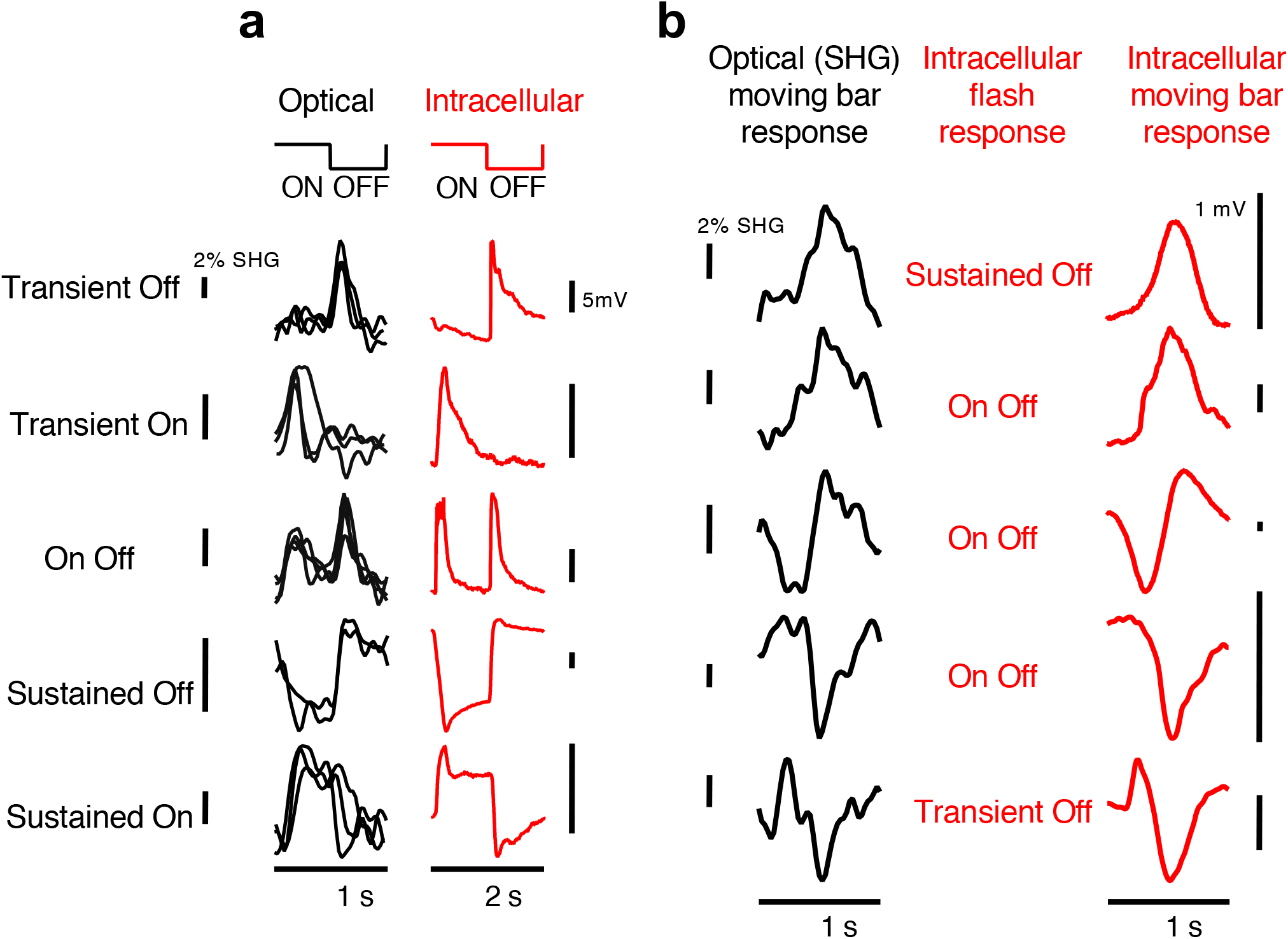
Comparison of intracellular and SHG responses from amacrine cells. (a) Left. SHG responses (black) from different types of amacrine cells to a uniform field flash of light at 1 Hz, each trace a different cell, showing representative examples for the five major types. Responses are the average of 140 trials. Right. Example intracellular recordings (red) recorded separately from different cells and retinas to a uniform field flash of light at 0.5 Hz. (b) Moving bar responses from individual amacrine cell SHG recordings (black) compared to similar single cell intracellular recordings from different preparations (red) classified by their intracellular flash response. All responses are to black bars moving against a bright background. SHG responses are classified as Off for the top two cells, Biphasic for the middle cell, and On for the bottom two cells.

**Figure S5.**
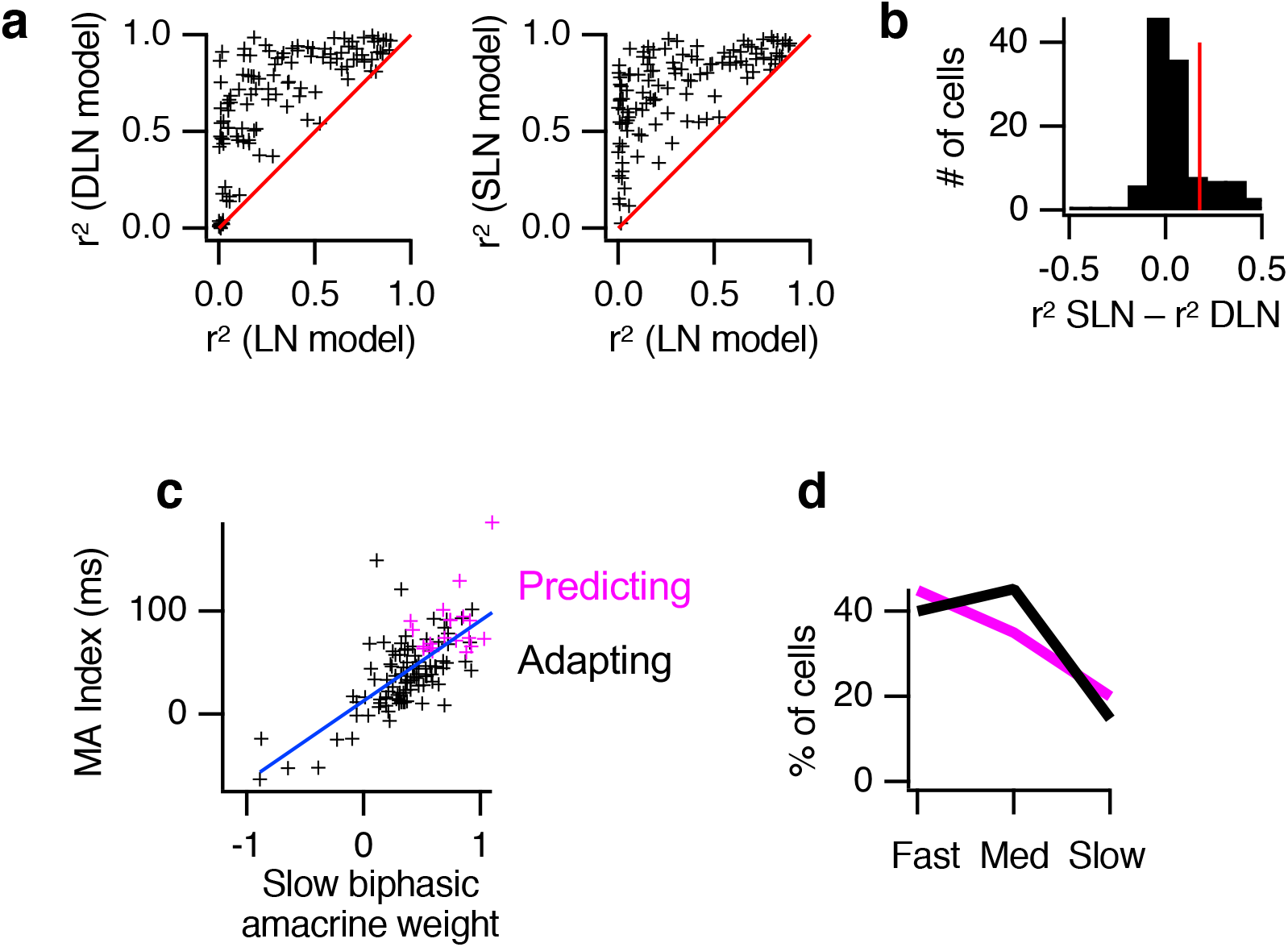
Model of amacrine contribution to motion anticipation. (a) Left. Scatterplot of squared correlation coefficient (r^2^) for the DLN model (divisive, average r^2^ = 0.66) compared to the LN model (average r^2^ = 0.32). Red line is the identity. Right Same for the SLN model (subtractive, average r^2^ = 0.72) compared to the LN model. (b) Histogram of difference in r^2^ for the SLN and DLN models (SNL -DLN). A k-means clustering with a value of 2 clusters (average Silhouette score = 0.89) divides the data into two classes, the Motion Adapting group (102 out of 122 cells) that are equally well fit by SLN and DLN models, and the Motion Predicting group where the SLN model exceeds the DLN model in accuracy (20 out of 122 cells). (c) Scatterplot comparing the motion anticipation index and the weights on the slow biphasic type of amacrine cells for the SLN model (red = adapting cells, black = predicting cells, r = 0.7, p[t2] < 3.9e-18). (d) For Motion Adapting and Motion Predicting cells, the fraction of cells that have different temporal filter speeds (fast, medium, slow). Note that values are computed and shown separately for Motion Adapting and Motion Predicting cells.

## Materials and Methods

### Multielectrode recording

Multielectrode array methods for extracellular recording and analysis were as described (13). The salamander retina was isolated intact, and the photoreceptor side was adhered by surface tension to a dialysis membrane attached to a plastic holder. The retina holder was then placed on a motorized manipulator and lowered onto a 60-electrode array (ThinMEA, Multichannel Systems) ganglion cell side down. In this way, a large piece of retina, up to its entirety, was placed on the array with minimal disturbance to the neural circuitry. The multielectrode array was patterned on a coverslip (∼180 µm thickness) allowing imaging from below through the array. The 10 µm diameter electrodes were separated by either 100 µm (low-density array) or 30 µm (high-density array). The array formed the bottom of a perfusion chamber, through which flows oxygenated Ringer solution buffered with bicarbonate. The full waveform of signals from the array of electrodes were digitized at 10 kHz and recorded on a computer.

### Custom two-photon laser scanning microscope

A two-photon laser-scanning microscope was constructed in an inverted configuration to simultaneously perform multielectrode array recordings. A simplified diagram of essential elements is shown in (Fig. 1a). Excitation was provided by a mode-locked Ti:sapphire (Tsunami, Spectra-Physics) laser operated at 970 nm. For SHG imaging, three main optical pathways were used: (1) The laser beam entered the microscope from the bottom and was focused on the retina by a x40 1.2 NA (Zeiss) objective. (2) The SHG signal was collected from above the retina through a x40 0.8 NA (Leica) objective. This signal at half the excitation wavelength (485nm) was reflected (Z488RDC dichroic, Chroma) into a PMT (H7422-P, GaAsP from Hamamatsu) through a filter (FF01-488/20, Semrock) and a laser-blocking filter (FF01-680/SP, Semrock). (3) The visual stimulus delivered from a CRT video monitor was reflected to the microscope (FF735 dichroic, Semrock) and focused on the retina through the upper 0.8 NA x40 objective.

The multielectrode array and the retina were mounted on a fixed stage. To image different areas of the retina, the inverted microscope, including the x-y scanning galvanometer mirrors (VM-500, GSI Lumonics) were moved on an x-y stage. Each stage moved along the entering optical axis of the laser, thus not altering laser alignment. The software used to control the mirrors and save the images was Scanimage (Pologruto et al., 2003).

### Imaging

Voltage sensitive fluorescent dyes have been used to measure the membrane potential both of subcellular structures such as dendrites and axons, and to record cell populations in neural networks (Homma et al., 2009). Second harmonic generation (SHG) imaging has allowed the recording of membrane potential from single cells (9), as well as many neurons in a brain slice (17). Unlike two-photon fluorescence, where two coincident photons are absorbed and then a single photon is emitted after a shift in wavelength, SHG is an energy conserving process, converting two photons into a single photon at exactly half the wavelength.

The SHG signal is produced from polar molecules arranged in an anisotropic structure. Styryl dyes such as FM 4-64 arranged in a membrane produce a SHG signal that is sensitive to membrane potential. As the voltage-dependent SHG signal is primarily transmitted through the tissue – not emitted in all directions as in fluorescence imaging – the transparent retina is well suited to this technique.

Because the SHG signal is highly dependent on the membrane voltage, intracellular membranes contribute virtually no background signal, unlike fluorescent voltage sensitive dyes (18). In addition, a primary dye used for SHG imaging, FM 4-64, is well tolerated by neural tissue for prolonged periods of time (> 1 hour), unlike many fast voltage sensitive dyes that have a significant pharmacological effect on neural responses (Homma et al., 2009). FM 4-64 is most often used as a fluorescent dye simply to label membranes in studies of synaptic vesicle recycling, and thus this dye is well tolerated by neurons and synapses (19).

In SHG imaging, the largest signal is scattered in the forward direction (Moreaux et al., 2000). Normally the collection objective would have a numerical aperture equal to or larger than the excitation objective to collect all the scattered light (Moreaux et al., 2000). Because of mechanical constraints, including the retina holder and micropipette, a long working distance necessitated a lower numerical aperture objective. Thus we underfilled the 1.2 NA laser excitation objective so that it effectively functioned as a 0.8 NA objective, allowing the 0.8 NA collection objective to capture all of the forward-scattered SHG signal. The average power at the sample was ∼10 mW which is comparable to the laser power used (20) for calcium imaging in salamander retina (*λ*=930nm, 2-30mW).

The styryl dye FM 4-64 was bath applied by immersing the isolated retina in 82 μM (100 *µ*g in 2ml) FM4-64 in oxygenated Ringer’s for about 1 hr prior to its placement on the multi-electrode array. When the dye is applied extracellularly, a depolarization of the membrane reduces the measured SHG signal (Sacconi et al., 2006). SHG responses are displayed so that depolarization is shown in the upward direction. In these studies we did not directly measure the relationship between SHG intensity change and voltage change for the dye FM4-64, although in previous studies this relationship ranges from 7.5 to 14 % / 100 mV (17, 21). As the range of amacrine cell responses we measure range from 1 to 10%, this puts the response in the expected range (tens of mV) for amacrine cells. Note however, that because we measure SHG measurements for relative sensitivity to the stimulus and correlation with ganglion cells, we don’t interpret the absolute amplitude of the voltage response.

The laser scanned an area of 164 × 73 μm (288 pixels × 128 pixels) bi-directionally in x (fast) and unidirectionally in y (slow) at a frame rate of 20 Hz (0.4 ms/line) about 55 μm above the multielectrode array in the amacrine cell layer, which is in the inner nuclear layer just proximal to the inner plexiform layer (22, 23). The visual stimulus was uncorrelated with laser scanning so that each time the laser sampled a point in space it was in a different part of the stimulus cycle.

The SHG signal was strongly dependent on the polarization angle of the laser compared to the orientation of the dye molecules in the membrane (24). Two laser polarization angles that were 90° apart produce an SHG signal from different parts of the soma (Fig 2b, S1a), showing that signals were SHG and not fluorescence. For physiological recordings, only one laser polarization was used (0°, green areas in Fig 2b).

A second PMT located below the retina was used to capture fluorescent images for histological comparison with simultaneously measured SHG images. This was done when no visual stimulus was present, as the red emission fluorescence of FM 4-64 overlaps the red visual stimulus. A stack of images spanning 120 µm with 1 *µ*m spacing was taken to compare SHG and fluorescence signals (Fig. S1b). Both fluorescence and SHG signals were brighter in the IPL, by virtue of high density of processes there. However, the ratio of SHG to fluorescence differed across retinal layers, and was lower in the IPL than in the INL (Fig S1c). This differing SHG/fluorescence ratio likely relates to the requirement for molecular asymmetry in generating the SHG signal (Moreaux et al., 2000). In the INL, it is expected that oriented membranes and less densely packed molecules and would reduce the likelihood that the distribution of dye molecules is symmetric within a small volume.

### Laser scanning as visual stimulus in the retina

It is known that laser scanning can function as a visual stimulus at 930 nm in the salamander retina (20), which is dominated by long wavelength cones. Even at our chosen wavelength of 970 nm, at slow scan rates (< 10 Hz) we observed ganglion responses synchronized to the laser as previously observed (20). To test whether a 20 Hz scan rate would modulate the spiking response of ganglion cells, we measured the power of Fourier transform (FFT) of the extracellular ganglion cell response, and compared the response at 20 Hz to the response for a 1 Hz visual stimulus. At a 20 Hz scan rate, for all cells the FFT power at the laser scanning frequency was less than 1% of the FFT power at the visual stimulus frequency. However, it is possible that there are subthreshold effects that cannot be measured from spiking activity. We therefore performed intracellular recordings and tested the effect of laser scanning by itself and in combination with visual stimulus.

Figure S2 shows the response of an Off-type amacrine cell recorded intracellularly. In response to a 10 Hz scanning laser centered over the cell’s receptive field, there was a measurable response of 1 – 2 mV, compared to a response of tens of mV to a 1 Hz uniform-field visual stimulus. However, the response to the laser at 20 Hz was at least 3000 fold smaller as measured using an FFT, and was not detectable (Fig. S2a). We then examined the response to a uniform field Gaussian flicker at low contrast (12%) with and without laser scanning at 10Hz (Fig. S2c). We computed a linear-nonlinear (LN) model consisting of a linear temporal filter representing the average response to a small flash of light and a nonlinear response function that captured any rectification or saturation of the response (Baccus & Meister, 2002). The temporal filters and nonlinearities (Fig. S1c) were nearly identical with and without the laser. We chose a 20 Hz scan rate for our standard experimental condition.

### Photoelectric artifact

The laser produced a photoelectric artifact on the titanium nitride electrodes if the imaged area was located directly over the array. This problem was avoided by imaging adjacent to the array (Fig 2a). Amacrine and ganglion cell receptive fields thus partially overlapped, and therefore the imaged amacrine cells were likely exposed to greater background illumination from the laser than were ganglion cells. Although this additional background illumination would result in a slightly different state of luminance adaptation and potentially lower contrast, intracellular recordings indicated that the laser did not reduce sensitivity for the visual stimulus, even when directly centered over the receptive field center (Fig. S2b).

### Visual Stimuli

Visual stimuli including uniform flashes, random checkerboards and moving bars were projected from a CRT monitor using Psychophysics Toolbox. To prevent the PMT from detecting the visual stimulus, only the red gun of the CRT was used and two long pass filters (XF3094, 610ALP, Omega Optical) inserted in the visual stimulus pathway restricted the wavelength of the visual stimulus to longer than 610 nm. The mean luminance was approximately 30 mW/m^2^ and the diameter of the stimulus was ∼ 1 mm on the retina. For experiments presenting a moving dark bar the background illumination was ∼60 mW/m^2^. Periodic stimuli were generally repeated 140 times (1Hz, 140 s) and the average response is shown. Random checkerboard stimuli were presented for 150 s. As mentioned previously, laser light (one-photon, see (20)) can stimulate photoreceptors, likewise for fluorescence and SHG signals. These effects are minimized two ways: first, by having a sufficiently high frame rate (∼20Hz) so that this illumination serves as background only and does not modulate cell responses in time, and second, to use a high background luminance for the visual stimulus so that photoreceptors adapted to this background will be relatively insensitive to these other light sources.

### Regions of Interest

All somas in the inner nuclear layer generated an SHG signal. The average response image across the entire recording was processed to enhance its contrast, and then thresholded to identify regions of interest with a large mean SHG signal. When the dye FM4-64 is bath applied (as opposed to picospritzing) there is an initial few seconds when the mean SHG signal decreases but thereafter it remains stable for the rest of the recording (∼150 s). The time-varying signal for each pixel was then binned relative to the stimulus. These anatomically based ROIs might include pixels from more than one cell, so principal components analysis was used to identify pixels that were responding differently from the first principle component and these pixels were rejected from the ROI. ROIs were accepted as belonging to a cell if they were larger than 20 pixels, had a SNR of greater than 2, and did not span the membrane of more than one cell. This yielded ROIs with visual responses that overlaid cell membranes. Multiple ROIs that appeared to belong to the same cell were grouped together as one cell.

### Signal to Noise Ratio

Although SHG is capable of very high temporal resolution (∼1 ms) (Dombeck et al., 2005), we generally binned image data relative to either the stimulus or spike times by increments of 25 ms to improve the signal to noise ratio. The average correlation with both the visual stimulus and with ganglion cell spikes was comprised primarily of slower variation, consistent with electrophysiological measurements (7, 25). Some recordings, however, showed a higher SNR, demonstrating that high temporal resolution (5 ms bins) is possible with SHG imaging given sufficient SNR or signal averaging (Fig. S3a).

We also examined whether responses were stable over time during our recordings (Fig S3b). Although the amplitude of the response in terms of fractional change in SHG was stable over 140 s, noise did appear to increase slightly over time, likely due to a decrease in the absolute amplitude of the SHG response due to bleaching. Because signals were always computed as a fraction of SHG at the current time, this change over time did not influence our results.

### Receptive fields

Spatial receptive fields and temporal filters were calculated by the standard method of reverse correlation with a visual stimulus consisting of binary squares (26), such that

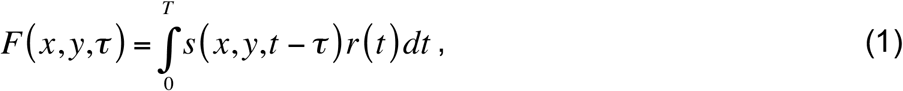

where *F* (*x, y,τ*) is the linear response filter at position (*x, y*) and delay *τ, s* (*x, y,t*) is the stimulus intensity at position (*x, y*) and time *t*, normalized to zero mean, *r* (*t*) is the firing rate of a cell, and *T* is the duration of the recording. The filter *F* (*x, y,τ*) was computed by correlating the visual stimulus to spike times for ganglion cells. A temporal filter was computed as the spatial average of *F*(). The stimulus *s* (*x, y,t*) was convolved with the filter *F* (*x, y,τ*) to obtain the linear prediction of the response. A static non-linearity was computed as the average relationship between the firing rate of the cell and the linear prediction.

### Modelling of motion anticipation by amacrine cells

#### LN model

The above mentioned receptive field filter and non-linearity were computed for each Off ganglion cell (n=115). A moving bar stimulus was convolved with the receptive field filter to obtain a linear prediction *L(t)*,

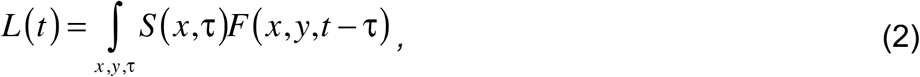

where *S* (*x*, τ) is the moving bar stimulus and *F* (*x, y*, τ) is the space-time receptive field filter computed from a white noise stimulus. The LN model consisted of the linear prediction and non-linearity being rescaled so that when the linear prediction is passed through the nonlinearity the peak firing rate of the LN model matched the actual peak firing rate. This rescaling represents the fact that maximal firing rate to a moving bar stimulus is generally much greater than to a white noise checkerboard stimulus,

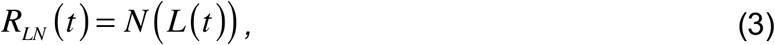

Where *N* () is the rescaled non-linearity computed from the white-noise stimulus and *R*_*LN*_ (*t*) is the firing rate of the LN model.

#### SLN model, subtractive inhibition and disinhibition

The filter of the LN model described above includes amacrine cell responses to a white noise checkerboard stimulus and thus it includes the linear component of the amacrine cell contribution to the ganglion cell receptive field. We wished to determine the possible nonlinear contribution that amacrine cells might have to change the response of the cell a moving bar stimulus. The general structure of this model was to optimize a weighted combination of amacrine cell type responses, which was used to modify the linear component of the LN model and then was passed through the same nonlinearity as the LN model to yield a firing rate output. Models were fit to minimize the mean squared error between the model output and one half of the data and then tested on the other half. All reported results are from this held out data.

Amacrine cell responses to the moving bar stimulus (n = 365) were clustered (by k-means) into three general categories as shown in Fig. 2f, (Off, Biphasic, and On) and then further divided the Biphasic class into Fast and Slow classes. We verified by F-test that the four parameter model (Off, fast Biphasic, slow Biphasic and On) was significantly better (p = 3.5e-11) than the three parameter model (Off, Biphasic, On) on held out data. Subdividing Off and On cell classes did not improve the model.

To account for transmission delays and spatial weighting connections between amacrine and ganglion cells, we used a temporal and spatial filter taken from (13), in which current was injected into sustained Off amacrine cells to measure a temporal filter and spatial spread of inhibition onto nearby Off ganglion cells

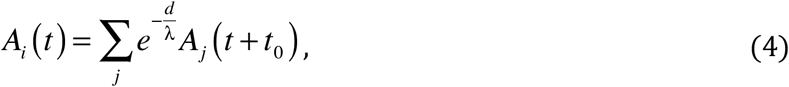

where *A*_*j*_ (*t* + *t*_0_) is an individual amacrine cell response to the moving bar whose response has been delayed by *t*_*0*_ (25 ms), which represents the delay of the amacrine cell transmission filter (13), *d* is the distance between amacrine cell *j* and a ganglion cell, *λ* (83 μm) is the space constant that describes the amacrine cell influence as a function of distance for a particular type of amacrine cell, (13) and *A*_*i*_ is the spatially weighted summed responses for one of the four types of amacrine cells *i*. The amacrine input *A*_*i*_ was computed separately for each ganglion cell because every ganglion cell will have different spatial weightings from the same amacrine cells. The spatially weighted summed amacrine responses *A*_*i*_ were then optimally weighted to minimize rms error between the model and data by *w*_*i*_ and summed:

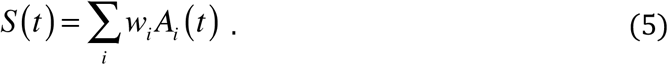

A single value of *w*_*i*_, which was allowed to be positive or negative, was applied to each of the four amacrine cell types, so this spatial weighting consisted of four parameters. The total subtractive amacrine input *S(t)* was subtracted from the linear prediction *L* from the LN model and passed through the same non-linearity to obtain the firing rate for this version of the model,

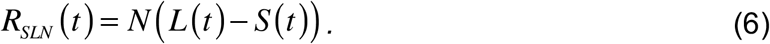

### The DLN model: divisive gain control

Alternatively, the linear prediction could be modified by dividing by amacrine responses. To avoid division by a small or negative number, the following formulation was used:

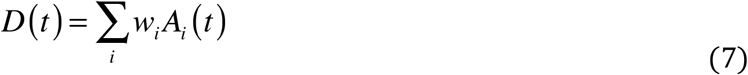

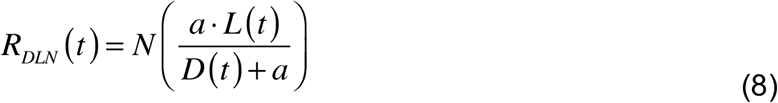

The constant *a* was a global free parameter fit to minimize population rms error. Both subtractive and divisive models were formulated so that when all the weights *w*_*i*_ are zero these models are equivalent to the LN model. The *A*_*i*_ terms were normalized in rms amplitude so that the weights *w*_*i*_ can be interpreted as the relative contribution of a particular type of amacrine cell.

It is important to point out the known limitations and simplifications of this model given what we know about the retina. Ideally there would be different space constants *λ* for each of the four types of amacrine cells, but this is unknown, and a limitation of the model. However, because nearby effects are considered and the moving bar is spatially localized in the direction of motion, it is expected that long-distance connections do not affect the response except in the direction of the extent of the bar, which is the second spatial dimension that does not play a role in the model. Likewise, transmission filters may vary between different pairs of amacrine and ganglion cells. Finally, our definition of amacrine cell classes comes from the response to a moving bar, and likely contains multiple types of amacrine cells.

### Model fitting

Because the fit of the model consisted only of four (SLN) or five (DLN) free parameters, models were fit by a grid search by calculating the total error for every value of weight in the stimulus space (−1.0 to +1.0) and choosing the solution with the lowest total error. The total error, *E*, consists of the RMS error between the data and either the divisive or subtractive model plus a quadratic penalty (*α*) on the sum of the weights:

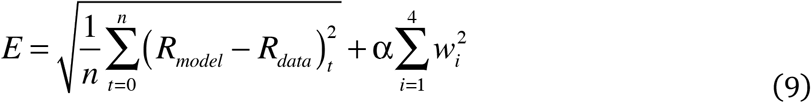

The value of *α* was determined by a 10% increase in RMS error compared to the RMS error when *α* = 0. This penalty on the weights reduces overfitting and drives the solution towards smaller values of weights which is relatively closer to the simplest model: the LN model. The value of *α* for the divisive model was 0.71 and the value for the subtractive model was 0.824.

